# Reinnervation of Muscle Targets Enhances the Separability of Motor Unit Signals Following Peripheral Nerve Transfers

**DOI:** 10.64898/2026.01.30.700058

**Authors:** Kiara N Quinn, Siyu Wang, Lang Qin, Alessandro Ascani Orsini, Kenzi Griffith, Rachana Suresh, Fausto Kang, Pierce L Perkins, Neha Joshi, Alexis L Lowe, Sami Tuffaha, Nitish V Thakor

## Abstract

After amputation, advanced prosthetic limbs offer a promising means of restoring motor function. However, state-of-the-art prostheses often rely on aggregate electromyogram (EMG) signals to decode motor intention, which limits their ability to replicate natural limb movements. Decomposing EMG signals into individual motor unit components has shown potential for more natural control, but distinguishing between individual units can be challenging when nearby signals overlap. This study demonstrates that muscle target reinnervation surgeries can naturally increase physical separation between motor unit signals, thereby mitigating this overlap. Reinnervation of individual motor units is evaluated in a rodent hindlimb model after direct nerve-to-muscle implantation. Histological and electrophysiological analyses reveal that *structural changes* following reinnervation surgery result in beneficial motor unit *signal changes*, particularly improving spatial separation between motor unit signals compared to those in intact muscle. This spatial separation contributed to fewer instances of complex, overlapping signals in reinnervated muscle recordings. Motor unit signals were leveraged to provide a proof-of-concept of precise control of a virtual prosthesis for the first time after direct nerve-to-muscle implantation surgery. These findings highlight the potential of reinnervated muscle targets as key biological interfaces that facilitate motor unit separation, reducing the burden on decomposition algorithms and improving prosthetic control.

## 1. Introduction

Following amputation, state-of-the-art robotic limbs offer the potential to restore motor function ^[1,2]^. However, even the most advanced prosthetic devices fall short of replicating the full range of capabilities, natural motion, and dexterity of the lost limb, leading to high rates of device abandonment ^[3]^. In an intact limb, precise movement control emerges from the coordinated activity of numerous motor units, which serve as the basic building blocks of the neuromuscular system ^[4]^. Each motor unit, defined as a single motor neuron and the muscle fibers it innervates, contains detailed information about intended movement dynamics. In contrast, control of artificial limbs often relies on the aggregate electromyogram (EMG) signal produced by the muscle, which provides an approximation of summed motor unit activity, but overlooks finer details ^[5,6]^. Therefore, current control strategies harness only a fraction of the available biological information about movement intention. To better replicate the natural precision of a wide range of movements in a robotic limb, recent research has shifted towards developing ways to reliably acquire individual motor unit signals to closely emulate the human motor system ^[7–9]^.

Advanced decomposition algorithms offer a promising approach for recording EMG signals with high-density electrode arrays and separating the mixed signal into its motor unit components ^[10,11]^. By identifying each motor unit’s unique characteristics, these blind source separation algorithms can sort units without any prior knowledge about the sources, similar to how a person can aurally distinguish between two different conversations happening in the same room without prior knowledge of who is talking or the topic of conversation ^[12]^. Despite considerable advancements, decomposition algorithms continue to face challenges in complex scenarios. For example, when multiple motor units near the same electrode contact are active simultaneously, their signals merge into a superimposed, complex waveform, making it difficult to distinguish between individual units ^[13]^. One approach to separate these overlapping signals involves algorithms like the peel-off method, which iteratively identifies and subtracts, or ‘peels-off,’ the most prominent motor unit signal from the composite waveform ^[14]^. However, this process is not as effective in cases of destructive interference, where two signals cancel each other out, or in cases of significant signal overlap ^[15]^. Alternatively, advancements in hardware, particularly high-density electrode technology, can enhance the isolation of individual signals. Theoretically, increasing electrode density increases the likelihood that each contact captures signals from fewer motor units, thereby reducing instances of signal superposition ^[16,17]^. Yet, in practice, the density of electrode contacts can be limited by physical constraints and cost considerations. These limitations highlight the need for innovative approaches to efficiently resolve superimposed waveforms.

To augment the existing software and hardware strategies above, we propose leveraging advanced surgical techniques, specifically muscle target reinnervation surgeries, to create natural physical separation between motor unit signals, thereby reducing signal overlap and complexity. Fundamentally, muscle target reinnervation surgeries help salvage nerve function after amputation by redirecting the axons regenerating from the severed nerve to a denervated muscle target **(Figure 1A)**. Once the target muscle is reinnervated by the nerve, it serves as a *biological amplifier* of neural signals, producing high-amplitude EMG signals that can facilitate intuitive prosthesis control ^[18–21]^. Herein, we aim to demonstrate that reinnervated muscle targets can also act as *biological separators*, resulting in larger physical separation between motor unit signals **(Figure 1B)**.

**Figure 1.**
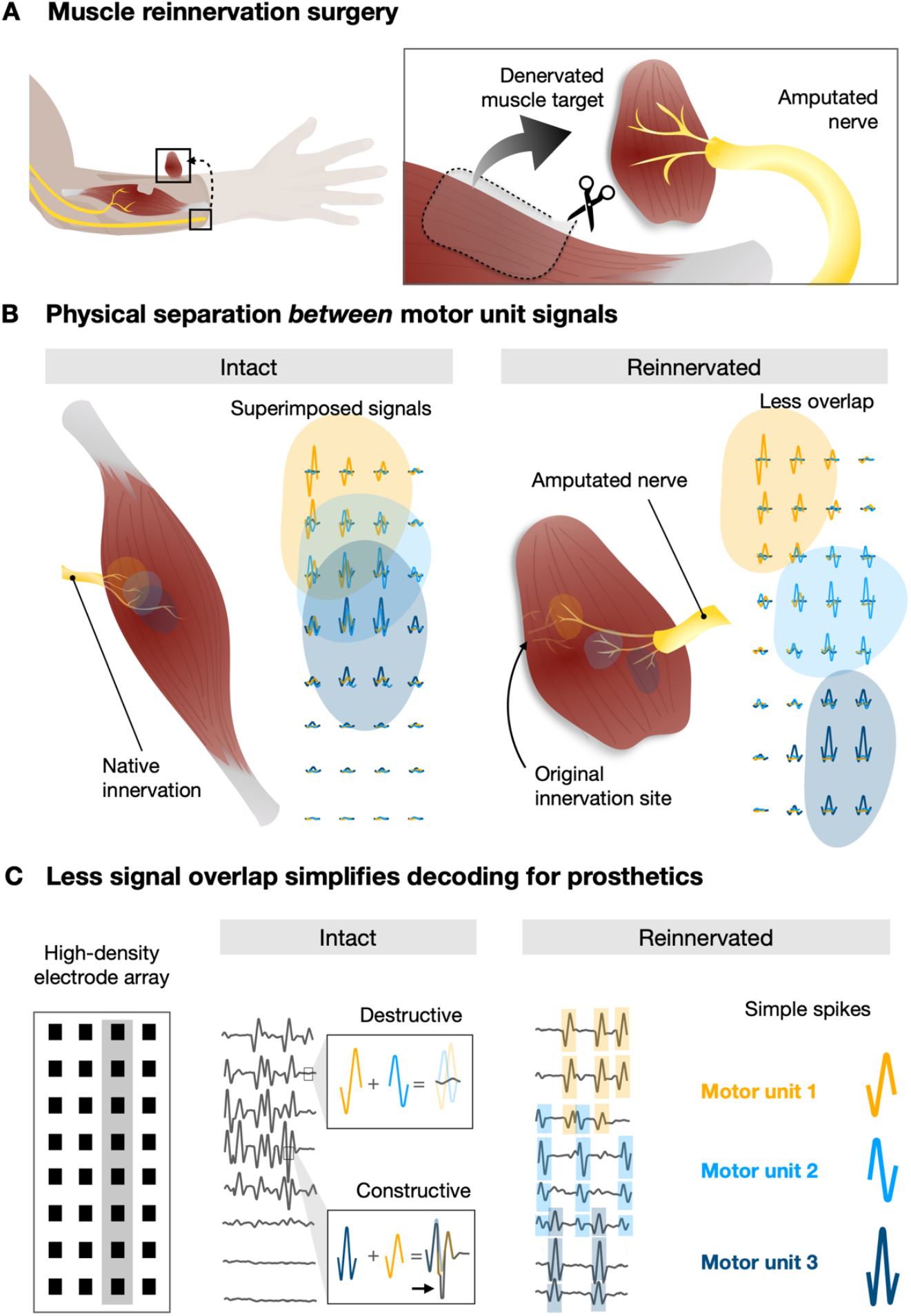
Overview of reinnervated muscle as a biological separator of motor unit signals. (**A**) Nerves severed during amputation can be surgically redirected to a new, denervated skeletal muscle target for reinnervation. (**B**) In intact muscle, motor unit signals can overlap spatially. In contrast, we hypothesize that structural changes that are inherent to reinnervation will lead to increased physical separation between motor unit signals. (**C**) We anticipate the increased spatial separation between motor units will lead to less signal overlap in reinnervated muscle compared to intact. Ultimately, we envision this enhancing the decomposition of motor units for improved decoding that will enable more natural prosthetic control that more closely emulates the fine motor control exhibited by the lost limb.

More specifically, we hypothesize that two structural changes inherent to the reinnervation process will contribute to less overlap between motor unit signals in reinnervated muscle targets compared to natively innervated, intact muscle **(Figure 1C)**. Firstly, during the reinnervation process, axons can either 1) re-establish connections, known as neuromuscular junctions (NMJs), that were previously occupied by the original nerve or 2) create new connections in areas where these NMJs are not natively found ^[22,23]^. We hypothesize that the creation of abnormally located connections can lead to more physical separation between NMJs, resulting in enhanced spatial separation between motor unit signals compared to intact muscle. Secondly, previous histological studies have demonstrated muscle fibers that belong to a single motor unit ‘clump’ together following reinnervation, contrasting with intact fibers that are evenly distributed throughout the muscle ^[24,25]^. We aim to demonstrate that this structural clumping translates to smaller, more concentrated motor unit signal territories that minimize the overlap between different signals.

In this study, we achieve reinnervation in a rodent model by implanting the amputated nerve directly into the muscle target via ‘direct nerve-to-muscle neurotization’ ^[26]^. We intentionally place the nerve opposite of the original innervation site to maximize the amount of space for ectopic NMJs to form as axons regrew toward the original nerve (Figure 1B). After reinnervation surgery, we first validate that motor units successfully reinnervate the muscle by investigating compound motor signals across the reinnervation period (15, 50 and 90 days post-reinnervation surgery). Second, we show histological evidence of abnormal NMJ formation. Next, we use high-density implantable electrodes to show motor unit signals in reinnervated muscle targets are more spatially separable compared to intact controls. Finally, we demonstrate a proof-of-concept of harnessing motor units for precise control of a virtual prosthesis. Our findings show that structural changes during reinnervation translate to changes in signal separability, thus showcasing the capability of muscle target reinnervation surgery to naturally create physical separation between individual motor units for prosthesis control. By increasing the spatial distribution of motor units, this approach can reduce overlap between motor unit signals, leading to fewer instances of superimposed signals, mitigating the burden on advanced decomposition strategies.

## 2. Results

### 2.1. Motor units reinnervate muscle targets over time after direct neurotization

Multiple variations of muscle target reinnervation surgeries are performed in the clinic, including targeted muscle reinnervation (TMR) ^[19,21]^, regenerative peripheral nerve interface (RPNI) ^[18]^, and vascularized denervated muscle target (VDMT) ^[27]^, the last of which is utilized in the present study. Although the main objective of these variations is the same—salvaging residual nerve function by providing a muscle to reinnervate—each achieves this goal differently. The cut nerve in TMR is connected to the nerve that originally innervates the muscle target, in an approach known as nerve-to-nerve ‘coaptation’ ^[19]^. In contrast, in RPNI and VDMT techniques, the residual nerve is connected directly to the muscle belly, also known as nerve-to-muscle ‘direct neurotization’ ^[26]^. Unlike nerve-to-nerve coaptation, direct neurotization offers the surgeon the flexibility to vary the location of the nerve. We opted for this because we intentionally wanted to maximize the amount of space for ectopic NMJs to form by implanting the amputated nerve farther from the original innervation site. We transected the distal tibial nerve just before it splits into the medial and lateral plantar nerves and then transferred it to a denervated soleus muscle target in the rodent hindlimb **(Figure 2)** ^[27]^. The transected nerve was connected to the muscle target on the side opposite the original innervation site. The most common clinical practice is to direct the nerve, when possible, into the innervation zone for optimal reinnervation, which has also been reported in research studies ^[19,28]^. Thus, it was critical that we first assessed the quality of reinnervation in our variation. Reinnervation was assessed in three groups at 15 days, 50 days, and 90 days post-reinnervation surgery (n=8 rats per group). Additionally, in the 90-day group, innervation of the intact soleus in the contralateral leg was evaluated, serving as a control (Figure 2). We electrically stimulated the sciatic nerve to evoke compound motor action potentials (CMAPs) from the muscle as shown in **Figure 3A-C** and Movie S1. By incrementally increasing the amplitude of the stimulation pulse (Table S1), we were able to estimate the number of functional motor units innervating the muscle using methods from ref. ^[29]^ (Figure S1,S2). As expected, it was observed that the estimated number of functional motor units that successfully reinnervated the muscle target gradually increased from 15 to 90 days post-surgery (**Figure 3D**, means and standard deviations reported in Table S2). Further, an increasing trend in maximum CMAP amplitude was noticed across the reinnervation period (**Figure 3E**, Table S2). Although the signal amplitude of the reinnervated muscle was still lower than the intact soleus (*p* = 0.0262), the number of motor units at 90 days post-reinnervation was not significantly different from the contralateral control muscle (*p* = 0.7489). These results demonstrate that even with nerve implantation opposite of the original innervation site, a sufficient number of motor units are available to serve as control sources for prosthesis control after reinnervation.

**Figure 2.**
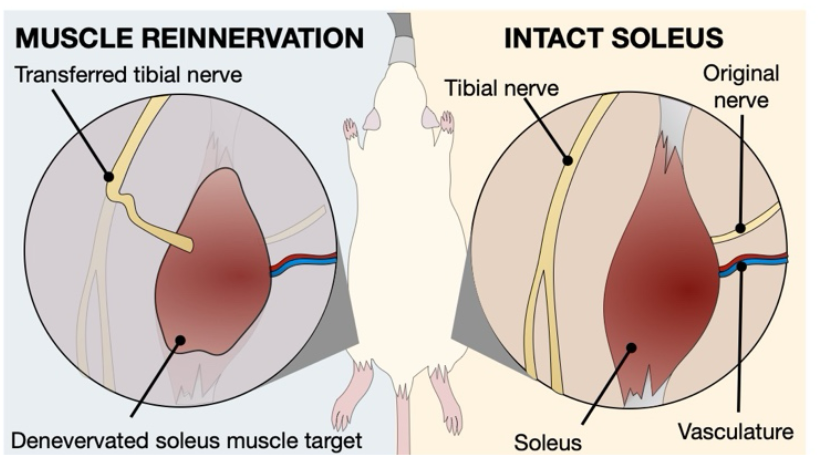
Rodent surgical model. The soleus muscle and distal tibial nerve were employed to replicate a clinically significant muscle reinnervation surgery. The tibial nerve was deliberately severed to mimic amputation and subsequently sutured to a denervated soleus muscle target for reinnervation. Importantly, the nerve was transferred on the opposite side of the original innervation site. Each rat underwent surgery on the left hindlimb while preserving the intact soleus on the right hindlimb as a control.

**Figure 3.**
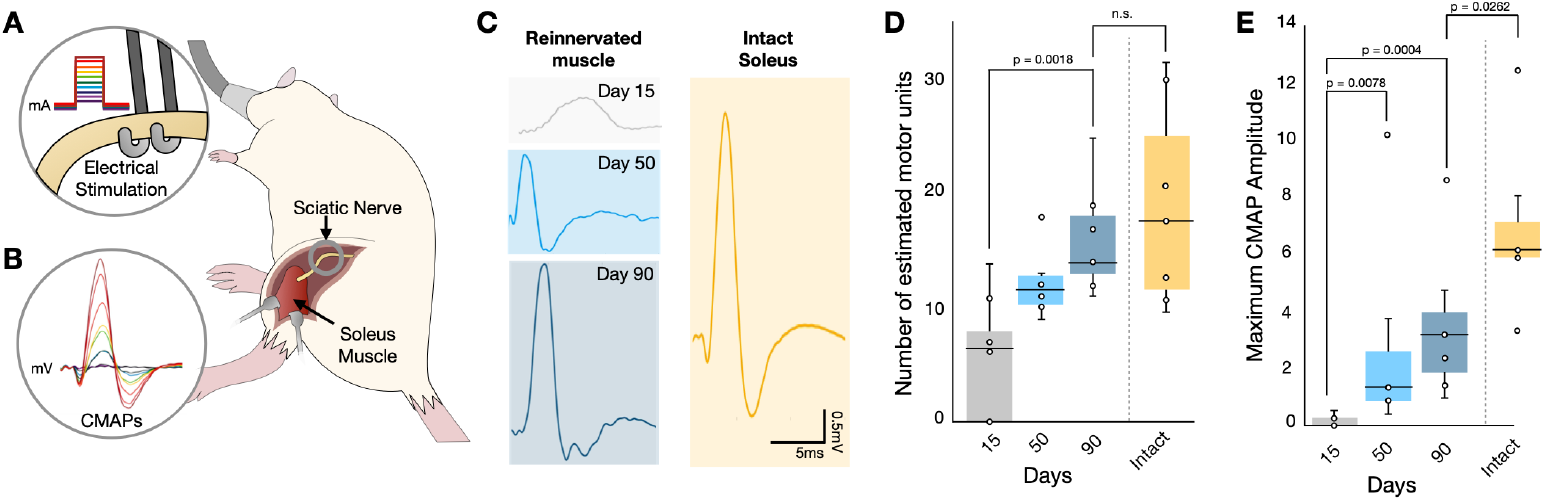
Motor unit number estimation and signal amplitude across reinnervation. **(A)** Fifteen, fifty, or ninety days post-reinnervation surgery, the sciatic nerve is electrically stimulated with incremental current pulses using a bipolar hook electrode. **(B)** Compound motor action potentials (CMAPs) are simultaneously recorded from the soleus muscle target using a fine needle electrode. **(C)** Representative CMAPs are shown for 15 days (gray), 50 days (blue), and 90 days (navy) post-reinnervation surgery in addition to the intact soleus (gold). **(D)** From the CMAPs recorded, we observed an increasing trend in the number of motor units reinnervating the muscle over time. Importantly, the number of reinnervated motor units is not significantly different from the intact muscle (p = 0.7489). **(E)** The maximum CMAP amplitude also increases over time, indicating that reinnervating axons are innervating more muscle fibers over time. The maximum CMAP amplitude at 90 days post reinnervation is still significantly lower than the intact soleus.

### 2.2. Spatial distribution of neuromuscular junctions changes after direct neurotization

We hypothesized that increased spatial separation between motor unit signals results from underlying structural changes that occur during reinnervation. NMJs serve as the physical connection between a peripheral nerve and its muscle target, facilitating the transmission of neural signals to the muscle. In healthy, intact muscle tissue, the spatial distribution of NMJs (i.e., location and density) is highly structured and conserved, varying according to the orientation of muscle fibers within specific muscle types ^[30]^. Following reinnervation surgery, the anticipated restructuring of NMJs was assessed using beta-III tubulin and alpha-bungarotoxin staining to identify axons and NMJ structures, respectively. Utilizing three-dimensional muscle tissue reconstruction, serial sections of muscle histology were stacked to enable visualization of NMJ distribution throughout reinnervated and intact muscle; averages of all the muscles within the intact and 90-day reinnervated group are shown in **Figure 4A-B**. In the top view of the intact muscle, few NMJs are in the proximal third region of the muscle (Figure 4A, Figure S3); in contrast, in reinnervated muscle tissue, a higher density of NMJs exists in the proximal third area in which the nerve was implanted during surgery (nerve implantation site is designated by a blue arrow in Figure 4A). Moreover, from the side view of the averaged muscles, there are an increased number of NMJs in the upper right quadrant (Figure 4B, Figure S3, Movie S2). To quantify the spatial separability of NMJs, the distances between NMJs on each section were modeled using a Rayleigh distribution. The spread of this distribution (α) offers insights into the dispersion of NMJs within the tissue sample. Higher spread implies wider dispersion of NMJs, whereas a lower α suggests a higher number of NMJs are in closer proximity to one another. Notably, NMJs in reinnervated muscle targets at 90 days post-surgery exhibited a wider distribution (α = 2.48 ± 0.85) compared to intact muscle (α = 1.54 ± 0.03, *p* = 0.0571, **Figure 4C**). Furthermore, we show evidence of clumping within motor units that aligns with the findings of previous studies ^[24,25]^; **Figure 4D** depicts NMJs from one motor unit clustered close together in space. This clumping pattern is hypothesized to contribute to smaller motor unit areas and minimal overlap between motor unit signals, as explored in the following section.

**Figure 4.**
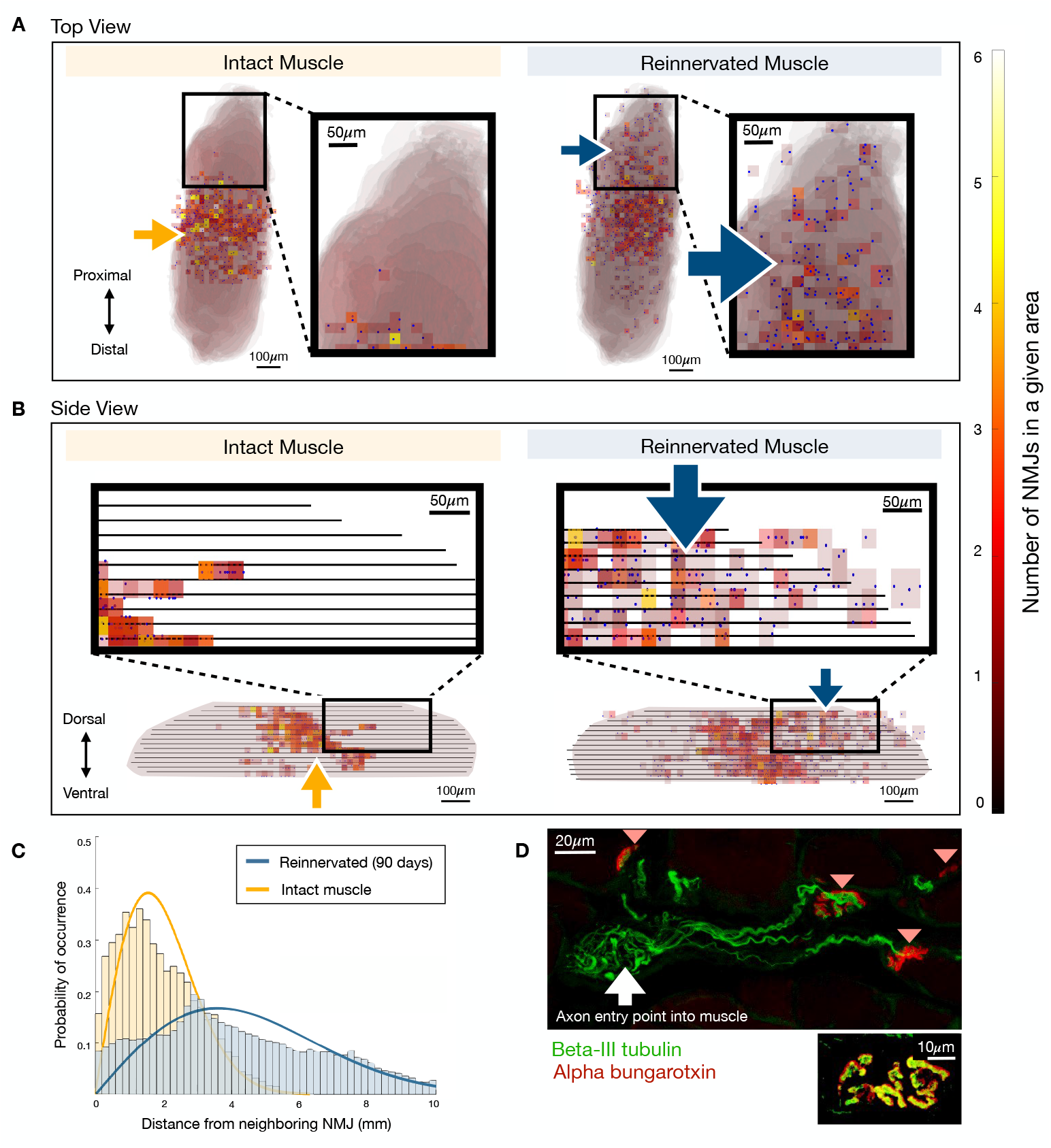
Neuromuscular junction distribution and structural changes leading to separability. Average NMJ distributions across all muscles are shown from the top and side view in (**A**) and (**B**), respectively. A yellow arrow designates the original innervation site, whereas the blue arrow designates the implantation site of the transferred nerve. Notably, there are a large number of NMJs near that transferred nerve in reinnervated muscle that are not normally present in intact muscle. (**C**) This formation of ectopic synapses leads to the wider distribution of NMJs in reinnervated muscle compared to intact muscle, leading to enhanced separability. (**D**) Clumping within a motor unit can occur when a single axon enters one point on the muscle and reinnervates NMJs nearby one another. Four reinnervated NMJs are labeled with a light red triangle. Axons and neuromuscular junctions (NMJs) are stained using beta-III tubulin (green) and alpha-bungarotoxin (red). A zoomed in view (bottom) shows large regions of yellow overlap, showing successful reinnervation.

### 2.3. Spatial separation between motor unit signals

To assess whether changes in NMJ spatial distribution manifested as changes in motor unit signal distribution, we employed high-density electrode recordings to create spatial maps of motor unit electrophysiology. More specifically, we utilized two 32-channel epimysial arrays per muscle ^[16]^ (Figure S4). To ensure consistent placement across muscles, we positioned two rows of electrodes above the nerve implantation site and evenly spaced the columns to cover the full width of the muscle. Individual motor unit signals were isolated from the summed activity recorded at each electrode contact to generate an interpolated spatial map of motor unit activity (**Figure 5A-D**). The spatial maps of reinnervated muscle (**Figure 5E**) exhibited fewer instances of overlapping motor unit territories (territories were defined herein as regions with >90% signal intensity) compared to those in intact muscle (Figure 5F and Figure S5). Importantly, increased distance between NMJs showed a strong positive correlation with increased distance between each motor unit signal’s center of mass (ρ = 0.82, *p* = 0.0235), demonstrating that structural changes in muscle physiology manifested as changes in motor unit signals (**Figure 5G**).

**Figure 5.**
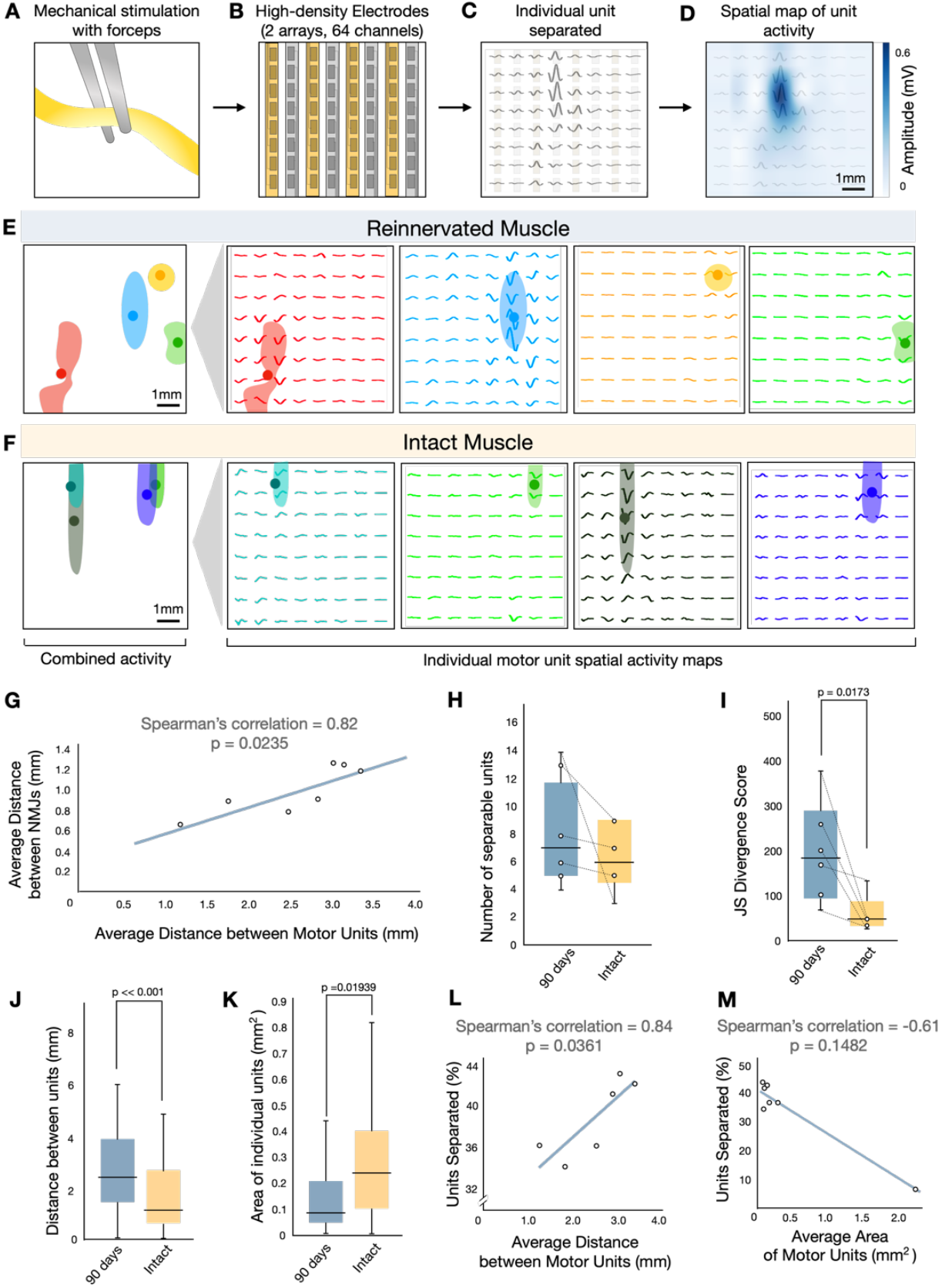
Spatial mapping of motor unit activity and functional separability. (**A**) The sciatic nerve is mechanically stimulated with forceps proximally to distally. **(B)** Motor units are recorded using two high-density electrode arrays. **(C-D)** Motor units are separated using Kilosort3 and heatmaps are generated using the root mean square (RMS) value of the average matched signals and cubic interpolation. Representative spatial maps of activity from four randomly selected motor units reveal less overlap between motor unit territories in **(E)** reinnervated compared to **(F)** intact muscle. **(G)** Increased average distance between NMJs is strongly correlated with increased average distance between motor unit signals. **(H)** There is not a statistically significant difference between the number of separable units in reinnervated and intact muscle, but within individual animals, there is a slight trend toward more separable units in reinnervated muscle compared to the contralateral leg. **(I)** Lower Jensen-Shannon divergence scores indicate more similarities between different motor unit territories (e.g., size, shape, location) within a given rat. These results demonstrate that reinnervated muscle has more distinct motor units than intact soleus, indicating increased separability. (**J**) Distance between motor units was higher and **(K)** area of individual territories was smaller in reinnervated muscle was higher compared to controls. **(L-M)** We show this increased average distance and decreased average area is contributes to the percentage of units that can be separated in reinnervated muscle.

The average number of separable motor units, conservatively defined as those with <70% overlap with another motor unit territory, was 7.88 ± 3.76 and 6.33 ± 2.42 in reinnervated muscle 90 days post-surgery and intact muscle, respectively (**Figure 5H**, *p* = 0.647). To quantify how distinguishable these units were from one another, we utilized Jensen-Shannon Divergence to assign a numerical score reflecting the dissimilarities between each motor unit’s spatial territory in each animal (e.g., size, shape, location of motor unit territory) ^[31]^. We observed that motor units at 90 days post-reinnervation were more dissimilar in size, shape, and location (198.38 ± 113.42) compared to units in intact controls (59.82 ± 42.81, *p* = 0.0173), which indicates a more distinct or separable spatial distribution (**Figure 5I**). We then further analyzed how much of this dissimilarity/separability was a result of *physical separation* between motor unit territories. It was observed that the distances between motor units’ center or masses are larger in reinnervated muscle at 90 days (2.69 ± 1.52 mm, n=349 units) compared to intact controls (1.65 ± 1.34 mm, n=116 units), emphasizing the enhanced physical separation between motor units (Figure 5J, *p* = 5.431 x 10^-11^). We show that the areas of individual motor unit territories were smaller in reinnervated muscle (0.13 ± 0.12 mm) compared to intact (0.35 ± 0.52 mm, Figure 5K, *p* = 0.0194). Smaller territories imply the signal is more concentrated in one area, which can contribute to less overlap between units. We performed a correlation analysis to determine the effect of physical separation between units and area of each unit territory on the number of units that can be separated (**Figure 5L-M**). As expected, average physical distance between units was found to have a strong positive correlation with the percentage of units that could be separated (ρ = 0.84, p = 0.0361); the average areas of unit territories were found to have a mild to moderate negative correlation with the percentage separable (ρ = -0.61, p = 0.1482).

Next, we further parsed out how the physical separation between unit territories affected the recorded signals, particularly the frequency of superimposed, complex signals and the ability of our sorting algorithm to distinguish between signals. Each spike detected in the recording was blindly rated as either simple or complex (**Figure 6A**). The percentage of complex spikes out of the total number of spikes in the recordings was higher in intact muscle (32.66 ± 4.18%) compared to reinnervated (20.87 ± 4.86%, *p* = 0.0006, **Figure 6B**). Correlation analysis revealed a higher percent of complex spikes was modestly associated with decreased average distances between motor units (**Figure 6C**, ρ = - 0.64, p = 0.1194) and increased average areas of individual units (**Figure 6D**, ρ = 0.5, p = 0.2532).

**Figure 6.**
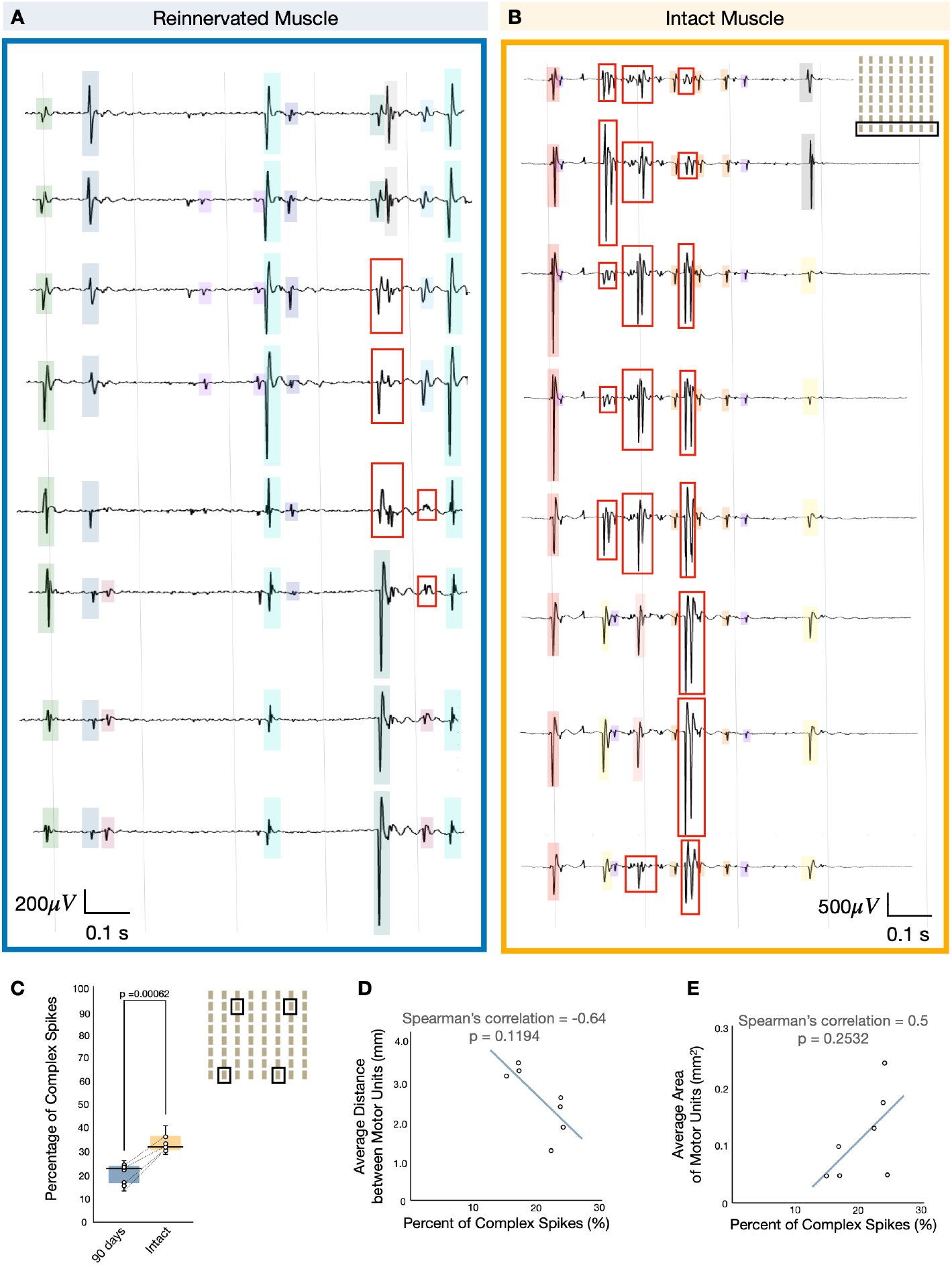
Signal complexity and superimposed spikes. Sample recordings from 8 electrodes are shown from reinnervated and intact muscle in (**A**) and (**B**), respectively. Complex waveforms are designated by a red box. (**C**) Notably, there is a lower percentage of complex spikes in recordings from reinnervated muscle compared to intact. The electrode array diagram shows the four channels (boxed) used in the analysis of percentage of complex spikes. (**D**) Correlation analysis reveals this reduction in signal complexity can be partly attributed to the increased distance between units and decreased area of individual territories.

### 2.4. Volitional motor unit activity can be used to predict movement intention after direct neurotization

In the experiments above, recordings (i.e., CMAPs and spatial maps) were achieved using artificial stimulation methods in fully anesthetized rodents. However, since the location of the nerve was varied compared to standard practice and this is the first investigation of motor unit reinnervation in a direct neurotization model, it was essential to validate our approach and determine whether volitional motor unit activity can be effectively translated into command signals for prosthetic movements. In other words, it is unimportant whether a large number of motor unit territories are separable if they fire indiscriminately, or all convey the same information. To address this question, the animal was placed under a light plane of anesthesia, such that volitional movement could be elicited by cutaneous stimulation. Rodents point their toes (plantar flexion) in response to cutaneous stimulation of the paw. The force produced during the plantar flexion was measured using a conductive rubber force sensor, serving as a proxy for movement intention (**Figure 7A**). A single MyoMatrix array was implanted percutaneously in an intramuscular configuration to record motor unit activity from a reinnervated muscle target (90 days post-surgery) during volitional plantar flexion (n=31 trials). Spatial maps of volitional motor unit activity are shown for two representative trials: one when the animal was producing low force (**Figure 7B**) and another producing high force (**Figure 7C**). Notably, motor units 1-6 were active over a larger area when higher force was produced compared to motor units 1-3 in a more concentrated area for lower force production (Figure 6B-C, Movie S3). This trend was consistent across other trials (Figure S6); a greater proportion of motor units were recruited during higher force outputs (46.18% ± 32.35%) compared to lower force outputs (13.34% ± 8.08%, **Figure 7D**, *p* = 0.0015). Additionally, the average firing rates of individual motor units were significantly higher during higher force outputs (7.56 ± 3.85) compared to lower force outputs (2.72 ± 1.22, Figure 7D, *p =* 0.027). Motor unit decomposition was performed using the same algorithm method as above. We then demonstrate volitional motor unit activity can be used to reconstruct the measured force curve (**Figure 7E, F**). The accuracy of the reconstruction was measured by demonstrating a high correlation and low mean squared error between the original and reconstructed curves (**Figure 7G**). This reconstructed signal was then used for precise control in a virtual prosthetic limb as shown in **Figure 7H** and Movie S4 to show the potential translational utility of reinnervated motor units. These results verify that even after reinnervation via direct neurotization, motor unit discharge patterns can be used to infer movement intention for prosthesis control.

**Figure 7.**
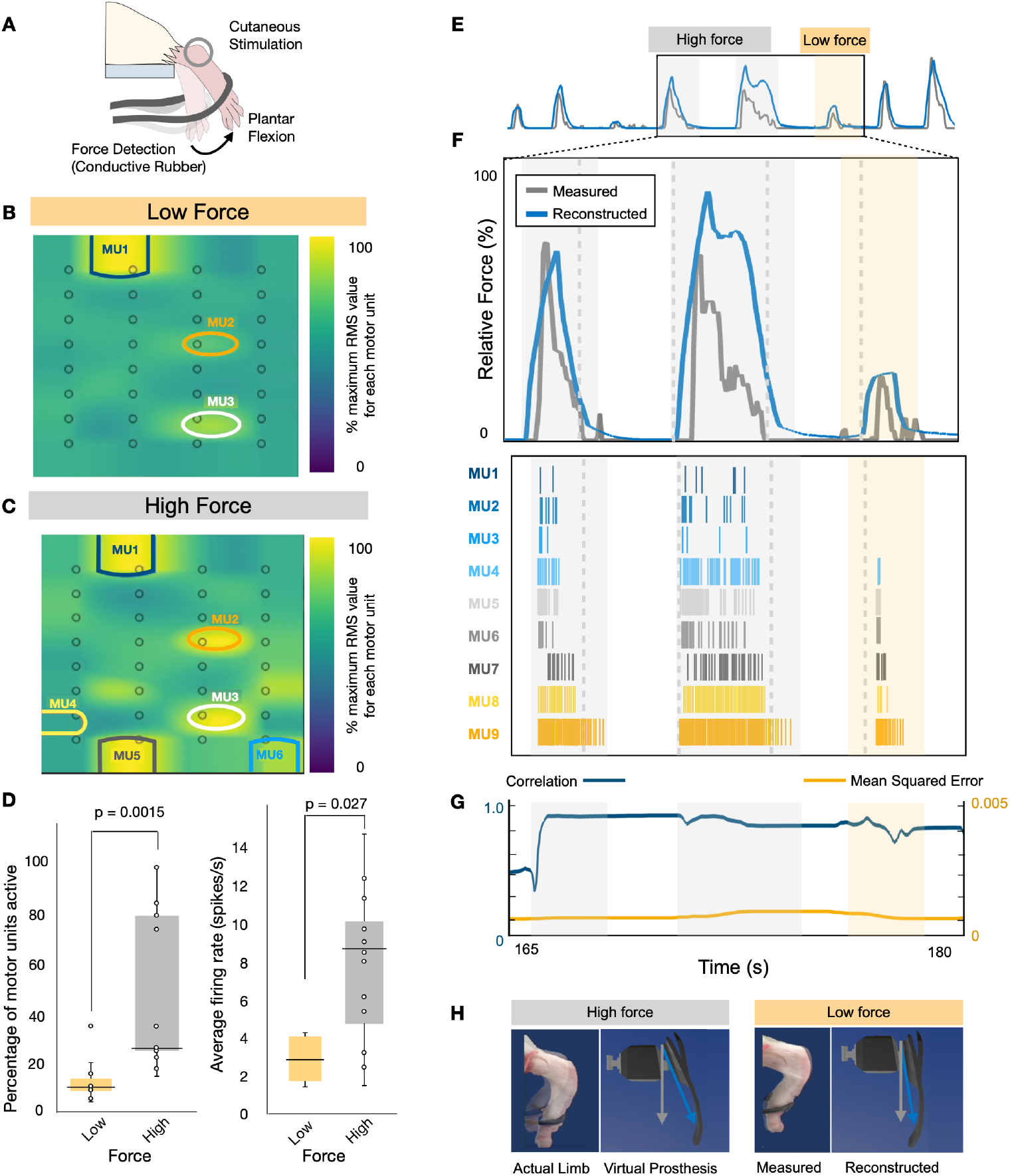
Motor unit activity can be used to reconstruct force production and control a virtual prosthesis. (**A**) To evoke volitional motor activity, cutaneous stimulation is applied to the paw of a restrained rat. This results in volitional plantar flexion of the ankle joint. The force produced during this motion is recorded using a conductive rubber force sensor. Motor unit activity is recorded by a 32-channel array implanted percutaneously. Representative heatmaps of spatial activity during a 0.05 second time window are shown for **(B)** low force (gold) and **(C)** high force (gray) production. **(D)** On average, higher force production during volitional movement is associated with a significantly higher percentage of recruited motor units and a higher average firing rate. (**E**) The motor unit activity pattern is robust enough to enable force reconstruction across all trials. Eight representative trials are depicted, including one low-force and two high-force scenarios shown in (**F**) the inset image. Motor unit raster plots across the three consecutive trials are shown. (**G**) The measured and reconstructed plots are highly correlated with a low mean squared error, emphasizing the utility and robust nature of the reinnervated motor unit activity. (**H**) Finally, motor unit activity was used to control the joint movement of a virtual prosthetic limb that closely mimics that actual movement of the animal.

## 3. Discussion

After amputation, advanced myoelectric prostheses can help restore motor function, but state-of-the-art devices still lack the precise control of a natural limb, leading to frequent abandonment ^[3]^. To address this discrepancy, recent research has leveraged high-density electrode arrays and decomposition algorithms to extract detailed motor unit activity for more natural control ^[7,8]^. A key challenge in the full realization of motor unit-based prosthesis control is that signal overlap leads to higher likelihood of superimposed waveforms, which can affect decomposition complexity. In this paper, we explored a novel use of an advanced surgical approach to enhance spatial separation of motor unit signals thus minimizing instances of complex, overlapping signals. Specifically, we utilized muscle target reinnervation surgeries, which have already gained traction as a therapeutic approach post-amputation, largely because they alleviate post-amputation pain ^[27]^ and can function as bio-amplifiers of residual nerve signals for prosthetic control ^[32]^.

For the first time, we illustrate the ability of reinnervated muscle targets to also serve as bio-separators of motor unit signals. We demonstrated that structural changes that occur after direct nerve-to-muscle neurotization, particularly increased distance between NMJs compared to intact controls (Figure 4A-C), manifest as increased distance between motor unit signals (Figure 5G). Further, we show the increased spatial separation leads to fewer superimposed, complex signals in raw recordings (Figure 6). Finally, we demonstrate a proof-of-concept of harnessing motor unit signals for virtual prosthesis control after direct neurotization surgery. Ultimately, this work has implications for facilitating the extraction of finer biological signals to enable more natural control of prosthetic limbs.

### 3.1. NMJs distributed across larger area after reinnervation via direct neurotization

The histological analyses in our study show that after reinnervation surgery NMJs were distributed across a larger area with increased distance between neighboring NMJs compared to intact muscle (Figure 4C). The average three-dimensional reconstructions of muscle sections for the intact and 90-day reinnervated group show evidence that this change in distribution presumably occurred due to formation of ectopic NMJs. We observed NMJs near the transferred nerve implantation site (designated by a blue arrow in Figure 5A-B) in reinnervated muscles that are not usually present in intact soleus muscles. This implies the implanted nerve created NMJs along the way as it grew from its implantation point toward the original nerve site (designated by a yellow arrow in Figure 5A-B). Doing so created a wider separation between NMJs throughout the reinnervated muscle target. This finding aligns with a previous study which shows axons can create ectopic synapses, especially in direct neurotization ^[22,23]^. We suspect that intentionally placing the nerve *across from* the original innervation, rather than nearby, contributed to the separation of NMJs as more area was available for ectopic synapses to be made.

Historically, most surgeons have aimed to transfer the nerve as close to the original innervation site of the target muscle, either by coapting the transferred nerve to the motor nerve supplying the target muscle or by implanting the severed nerve ending into the target muscle directly adjacent to the original nerve ^[19,28]^. This approach is intended to enhance the likelihood of successful reinnervation, as studies have shown that regenerating axons preferentially follow intramuscular neural tubes or “pathways” near the original innervation site to guide their growth ^[22]^. Importantly, we demonstrate that even in direct nerve-to-muscle neurotization when the nerve is placed far from the original neural tubes, the number of motor units following reinnervation of muscle targets was not significantly different from intact control muscles (Figure 3D). This finding is crucial as it suggests that in this surgical variation, patients could still retain an adequate number of healthy, reinnervating axons.

### 3.2. Enhanced spatial separation between motor unit signals in reinnervated muscle

One key novelty of this study is that it investigates how structural changes in NMJ distribution impact motor unit signals. Following reinnervation surgery, spatial mapping of activity reveals that individual motor unit signals exhibited less overlap compared to those in intact muscles (Figure 5E-F, Figure S5). Our analyses suggest that decreased overlap in territories can be attributed to the fact that motor unit signals in muscle targets are 1) more spatially separated and 2) smaller in area.

#### Distance between units

Our results show that the distances between motor unit signals were larger in reinnervated muscle compared to intact (Figure 5J), indicating that reinnervation surgery resulted in more spatially separated motor unit territories. Furthermore, we show a strong positive correlation between the average distances between motor units and the average distances between NMJs (Figure 5G). This supports our hypothesis that the structural changes following reinnervation are directly contributing to the changes in signal patterns. The increased distance between motor units also appears to positively affect the percentage of separable units (Figure 5L).

#### Area of individual units

Our results show the average areas of motor unit signal territories are smaller in reinnervated muscle compared to intact (Figure 5K). This aligns with previous histological studies that have found that motor units are fairly even in distribution throughout the muscle in intact muscles, such that in a muscle cross-section, fibers from different units appear in a mosaic pattern ^[24,33,34]^, whereas after reinnervation muscle fibers within a given motor unit tend to clump together ^[24,33,34]^. Our results illustrate how this clumping effect seen in histology can manifest as motor unit signal territories with smaller areas. Territories of smaller size could lead to easier decomposition for prosthesis control as the clumping of fibers from a single unit makes it easier to isolate that unit from others, supported by the moderate negative correlation between average motor unit area and the percentage of separable units (Figure 5M).

### 3.3. Reduced signal complexity in reinnervated muscle recordings

Ultimately, for prosthesis control applications, a major focus of this paper was to minimize signal complexity to enhance decomposition. Therefore, an important question to address was whether decreased overlap between motor units in reinnervated muscle would translate to fewer instances of superimposed, complex spikes. Our results showed the percentage of complex spikes is higher in intact muscle than in reinnervated muscle (Figure 6A-B). Correlation analyses suggest that this reduction in superimposed signals may be partly due to increased distance between motor units and decreased size of individual motor unit signals (Figure 6C-D). However, a limitation of this study is the small sample size, which may affect the reliability of our correlation analysis. Nevertheless, these findings suggest that patients who undergo muscle reinnervation surgery via direct neurotization could potentially benefit, as these effects could improve prosthesis control by simplifying the extraction of detailed neural signals.

### 3.4. Demonstration of motor unit-based prosthesis control after direct neurotization

#### To our knowledge, the current study represents the first comprehensive investigation of reinnervation at the motor unit level in a direct nerve-to-muscle neurotization model

Understanding motor unit reinnervation across different muscle reinnervation procedures can provide valuable insight into variations in patient outcomes and inform treatment strategies. We utilized force generation as our evaluation metric because, in the healthy neuromuscular system, force produced is a direct result of motor unit activity (i.e., recruitment and firing rate). During volitional movement in animals that underwent direct neurotization surgery, a greater percentage of motor units were recruited during trials where the animal generated high force compared to those with lower force output (Figure 7B-D). Additionally, individual motor units exhibited higher average firing rates in high-force trials (Figure 7D). By leveraging motor unit firing rates, we reconstructed force output and demonstrated its application in controlling a virtual prosthesis (Figure 7E-H). Together, these results demonstrated that even after direct neurotization, motor units still hold potential for decoding motor intention, specifically force generation. Future studies are necessary to further explore the full utility of motor unit control with reinnervated muscle targets in freely behaving animal models.

## 4. Conclusion

We have shown for the first time that structural changes in reinnervated muscle targets remarkably result in motor unit signals that are clumped and distributed in a larger space. This spatial separability of motor units can enhance the decomposition of motor units. These findings highlight reinnervated muscle targets as a key biological interface for harnessing motor unit signals for more natural prosthesis control that closely emulates the motor control strategies employed by the healthy nervous system.

## 5. Experimental Section/Methods

### Experimental Design

The broad objective of this study was to investigate motor unit reinnervation of skeletal muscle targets in a rodent hindlimb model. More specifically, the experiments presented herein were designed to analyze the spatial reorganization of motor units after muscle target reinnervation surgery. We aimed to show structural changes in NMJ distribution result in increased physical separation between motor unit signals. Twenty-six 8-week-old male Lewis rats underwent muscle target reinnervation surgery via direct nerve-to-muscle neurotization of denervated muscle targets on the left hindlimb. After the surgery, histological and electrophysiological data was collected to assess motor unit reinnervation (e.g., the spatial distribution of NMJs and motor unit signals in the muscle). This was a randomized, controlled study. All procedures were approved by the Johns Hopkins Animal Care and Use Committee.

### Surgical model

In this study, we modeled a clinically relevant muscle target reinnervation surgery (i.e., VDMT surgery) in the rodent hindlimb. In this study, we chose to model vascularized muscle target surgery because retaining a vascular supply can support larger muscle constructs and thus increase the number of available motor endplates for a larger nerve to reinnervate within a single muscle target ^[35,36]^.

The left hindlimb was sterilized and an approximately 3cm incision is made in the skin and biceps femoris muscle to expose the tibial nerve and soleus muscle. To mimic denervation due to amputation, we transected the branch of the distal tibial nerve 0.5 mm before it bifurcates into the lateral and medial plantar nerves using a surgical microscope. Then, a new denervated muscle target for the cut nerve to reinnervate was created from the soleus muscle. Specifically, the tendon attachments and original nerve supply were transected, leaving a denervated muscle flap of the soleus with only the blood supply left intact. A ‘pocket’ in the epimysium was made on the opposite side of the muscle respective to the original nerve’s innervation site. The implantation site was standardized across animals by using a fascial line on the soleus muscle as a reference point. The donor nerve (i.e., distal tibial nerve branch) was then implanted into the pocket and sutured to the VDMT using a 9–0 braided silk suture. The incision was closed using uninterrupted 4-0 sutures.

There are a few differences worth noting between clinical practice and our surgical model. First, redundant nerves and muscles are purposefully left intact such that rodent intent can be determined by observing the manifested movements performed by the animal. Secondly, in humans, a small flap of muscle is denervated. However, rodent muscles are inherently smaller and contain fewer NMJs than human muscle tissue. Thus, to ensure there are enough available motor endplates for the nerve to reinnervate, we use the entire soleus muscle. The muscle is de-inserted and de-originated to model a “flap.” Finally, surgeons will often wrap the muscle target around the nerve, creating what is colloquially known as a muscle ‘taco’ or ‘burrito.’ Here, we maintained the normal linearity of the muscle to preserve the spatial orientation/architecture of the muscle for histological analyses. A previous study noted this linear approach can improve reinnervation and pain outcomes in devascularized grafts ^[37]^.

### Motor unit number estimation

For each rodent (n=24), CMAPs were recorded from the left soleus muscle target to assess the functional innervation of the muscle. In the 90-day cohort, recordings were also performed on the contralateral intact soleus muscle as a control. CMAPs were utilized to estimate the number of functional motor units using methods adapted from ^[29]^. Briefly, to calculate the motor unit number estimation (MUNE), incremental steps of current stimulation amplitude were used to stimulate the sciatic nerve and recruit increasing numbers of motor units until saturation was reached (i.e., maximum CMAP amplitude). Stimulation (10 anodic square wave pulses at 10 Hz, 50μs pulse width) was achieved using a bipolar hook electrode (Natus Neurology Incorporated) while muscle responses were recorded using disposable 0.4mm diameter needle electrodes (Rhythmlink). Current amplitudes and current steps are detailed in Table S1. The average single motor unit action potential size is determined by taking the average amplitude of the first 10 evoked CMAPs. The estimated number of total motor units is then calculated by dividing the maximum CMAP amplitude by the average single motor unit action potential size (Figure S1). To minimize error from cross talk, all other branches from the sciatic nerve are severed before recording CMAPs (Figure S2, Movie S1).

### Spatial mapping of motor unit activity

Following CMAP collection in the 90-day group (n=8), two high-density electrode arrays (MyoMatrix arrays) were placed on the epimysium along the muscle fibers near the location of the nerve (each electrode array is 4 rows and 8 columns, totaling 32 channels; together, they create an array of 8 rows and 8 columns, totaling 64 channels) (Figure 5B) ^[16]^. To maintain consistency between animals, the arrays were placed such that two rows of channels were positioned proximal to the nerve implantation site (Figure S4). The sciatic nerve (which branches into the distal tibial nerve) was then mechanically stimulated (squeezed with forceps) proximally to distally for 0.5 seconds approximately 60 times (5 times at each location with incremental forces applied). Mechanical stimulation (as opposed to electrical stimulation) was used to ensure the artificially evoked motor unit activity had differences in both spatial and *temporal* characteristics, allowing for simplified and more accurate motor unit separation using Kilosort3. This stimulation strategy has been used in previous anesthetized animal studies of motor unit characteristics ^[28,38]^. The number (60 times) and duration (0.5 seconds each) of trials was determined to be sufficient to generate enough motor unit action potential trials for accurate decoding and separation. Data from the MyoMatrix arrays was recorded using the Intan RHD recording system (RHD2000, Intan Technologies) with a 20kHz sampling rate. Any channels with an impedance value > 50 kohms were excluded. Data was recorded bilaterally, with the starting side determined randomly. The recorded data from both intact and reinnervated muscle was then processed in two steps: 1) motor unit sorting and 2) motor unit visualization. For motor unit sorting, each motor unit is assumed to have a unique waveform. Using the Kilosort3 algorithm (parameters specified in Table S3), we can separate multiple motor unit ‘template’ waveforms, as well as the corresponding matched signals from the raw signal. The motor unit visualization step then involves using the matched signals for each motor unit to calculate the average waveform on each electrode channel. The root mean square (RMS) value from the average waveform is calculated to obtain the power intensity from each channel. Then, we apply a cubic interpolation to make a higher resolution 2D heat mapping image (Figure 5C-D). We quantified the separability of motor unit spatial territories by assigning a Jensen-Shannon Divergence numerical score to the motor unit territories in each muscle. Specifically, a higher divergence score denotes larger amounts of dissimilarities between motor units, indicating higher separability. We defined the area of each motor unit territory to be any region with an RMS value >90%. Moreover, we also defined each territory as a single point (center of mass of the area) to determine the physical distance between each pair of motor unit territories.

### Neuromuscular junction labeling and 3D reconstruction of serial sections

After completing electrophysiology recordings in the 90-day group, muscle tissue from both the experimental and control side was harvested and fixed with 4% paraformaldehyde for histological analysis (n = 8 rats). After a 24-hour fixation period, the tissue samples were cryoprotected using 15%, followed by 30% sucrose in phosphate buffer solutions (PBS) for 48 hours each. Tissue was then frozen in optimal cutting temperature compound, sectioned longitudinally along the fiber direction (20 μm thickness), and stained for beta-III tubulin (1:800 in PBS and 10% goat serum, Invitrogen) and alpha-bungarotoxin (1:800 in PBS and 10% goat serum, Invitrogen) to identify axons and NMJs. Every eighth section (160μm) was imaged using Nikon Eclipse Ti and analyzed using FIJI software. The number of NMJs was counted and the x and y locations of each NMJ were manually labeled. To quantify the spatial separability of NMJs, the distributions of NMJs on each section (i.e., x and y locations) were modeled using a Rayleigh distribution. Specifically, we calculated the distances between each pair of NMJs and created a histogram to show the frequency of each distance. Furthermore, a 3D physical structure reconstruction was made by stacking the resulting 2D images of serial sections using a custom MATLAB script described in detail in ^[39]^. To visualize average trends across muscles in each group, the 3D reconstructions of each muscle were aligned using their center point and tibial nerve implantation site as a reference. The area of the average 3D reconstruction was divided into small voxels of area 8 mm^3^. Within each voxel, the number of NMJs which exist within the voxel boundary was calculated (represented by a color scale) and the average location was labeled with a blue marker.

### Volitional motor unit activity, force reconstruction, and virtual prosthesis control

To record volitional motor unit activity, a single high-density MyoMatrix electrode array (4 by 8, 32 channels) was implanted percutaneously in rodents who had previously undergone VDMT surgery (90 days post-reinnervation). For the electrode implantation stage, the rodent was anesthetized using 2-3% isoflurane in a 1:1 oxygen-nitrogen gas mixture at a flow rate of 1L/min. The high-density array was implanted along the muscle fibers, uniformly across the soleus ^[16]^. To minimize cross talk, we placed the electrodes intramuscularly, with contacts facing the anterior compartment, isolating plantar flexion. To minimize electrode drift during motion, the electrode was sutured firmly within the muscle using 9-0 braided silk sutures. Once the electrode was successfully implanted, the incision site was closed, and the isoflurane dosage was lowered gradually to 1-1.5% to place the rodent under a light plane of anesthesia. The rodent’s left foot was placed on a conductive rubber stretch sensor (Adafruit Industries); changes in sensor length were monitored using an Arduino UNO, thus measuring the force generated. Volitional plantar flexion of the left ankle was evoked by applying light cutaneous stimulation on the right paw, resulting in a similar movement to the cross-extensor reflex. The approximate duration of cutaneous stimulation applied was kept consistent between trials. The strength of plantar flexion was volitional and thus randomly varied from trial to trial. The force measurement and the EMG signal were recorded simultaneously. A total of 31 trials were performed across two rats. Each force measurement was classified as either ‘high force’ or ‘low force’ based on whether it was ≥ or < the median force, respectively. It should be noted that this force is not the direct result of the soleus muscle target contracting; since it is a muscle target without tendons it is incapable of actuating movement of the joint.

Instead, we use the general foot movement as an indicator of intent because redundant muscles, such as the gastrocnemius, still control plantar flexion. Motor units were sorted as described in the ‘Spatial mapping of motor unit activity’ section. For each plantar flexion duration, the number of sorted motor units was counted. For each individual motor unit during the duration, the average firing rate was calculated. Additionally, a heatmap of combined motor unit activity (Movie S3) was generated by computing spatial mapping for all active motor units within a 0.05 sec window (0.01 sec step size). The overall firing rates of all motor units in each time window were utilized to construct a prediction of the distance the left paw was displaced (Figure 6E-F). More specifically, within each sliding window, the sum of all motor unit firing rates was divided by the sliding window size. The resulting overall firing rate curve was then normalized against its maximum point to produce a relative force (%) across different trials and animals. The predicted relative force output was used to control the movement of a virtual prosthesis in Blender 3D animation software using the built-in Python API (Figure 6H, Movie S4). In the virtual prosthesis, visualizing force production was challenging. Therefore, we assumed a constant and linear spring behavior for the conductive rubber stretch sensor. Using this assumption, relative force produced was proportional to the relative paw displacement based on Hooke’s Law, allowing for the control of virtual prosthesis displacement.

### Statistical Analysis

All statistical analyses were performed in RStudio. Results were reported as ‘mean’ ± ‘standard deviation.’ To determine whether each data set was normally distributed, the Shapiro-Wilk test was utilized. Non-parametric tests were performed when normal distribution and homogeneity of variances could not be assumed. Specifically, for analyses in which three different groups were measured (e.g., comparisons across 15-, 50-, and 90-day cohorts), the Kruskal Wallis was used to test differences between treated groups. If significance was found, a post-hoc Dunn’s test with Bonferroni correction was performed to determine which pairwise comparisons differed significantly. For 90-day rats, the untreated and treated sides were compared (right and left, respectively). Although these measurements were from the same subject, a paired test was not performed because 1) there can still be high variation within a single subject due to surgical variance, and 2) some values were missing. The Wilcoxon rank sum test was performed to determine any difference between the two groups. For correlation analysis between variables, Spearman’s correlation was calculated as it is more robust to outliers in small sample sizes and non-normal data compared to other methods.

## Acknowledgments

The authors would like to thank the members of the Center for Advanced Motor BioEngineering Research (CAMBER) for providing Myomatrix high-density electrode arrays for this study. We also express our gratitude to Deok Ho Kim’s lab at Johns Hopkins University for providing us with access to the Nikon microscope (Nikon Elipse Ti) and generously contributing their time and expertise through comprehensive training sessions.

## Data availability statement

The data that support the findings of this study are available from the corresponding author upon reasonable request.

## Ethics approval statement

All procedures were approved by the Johns Hopkins Animal Care and Use Committee.

