## Supporting Information for "Reinnervation of Muscle Targets Enhances the Separability of Motor Unit Signals Following Peripheral Nerve Transfers"

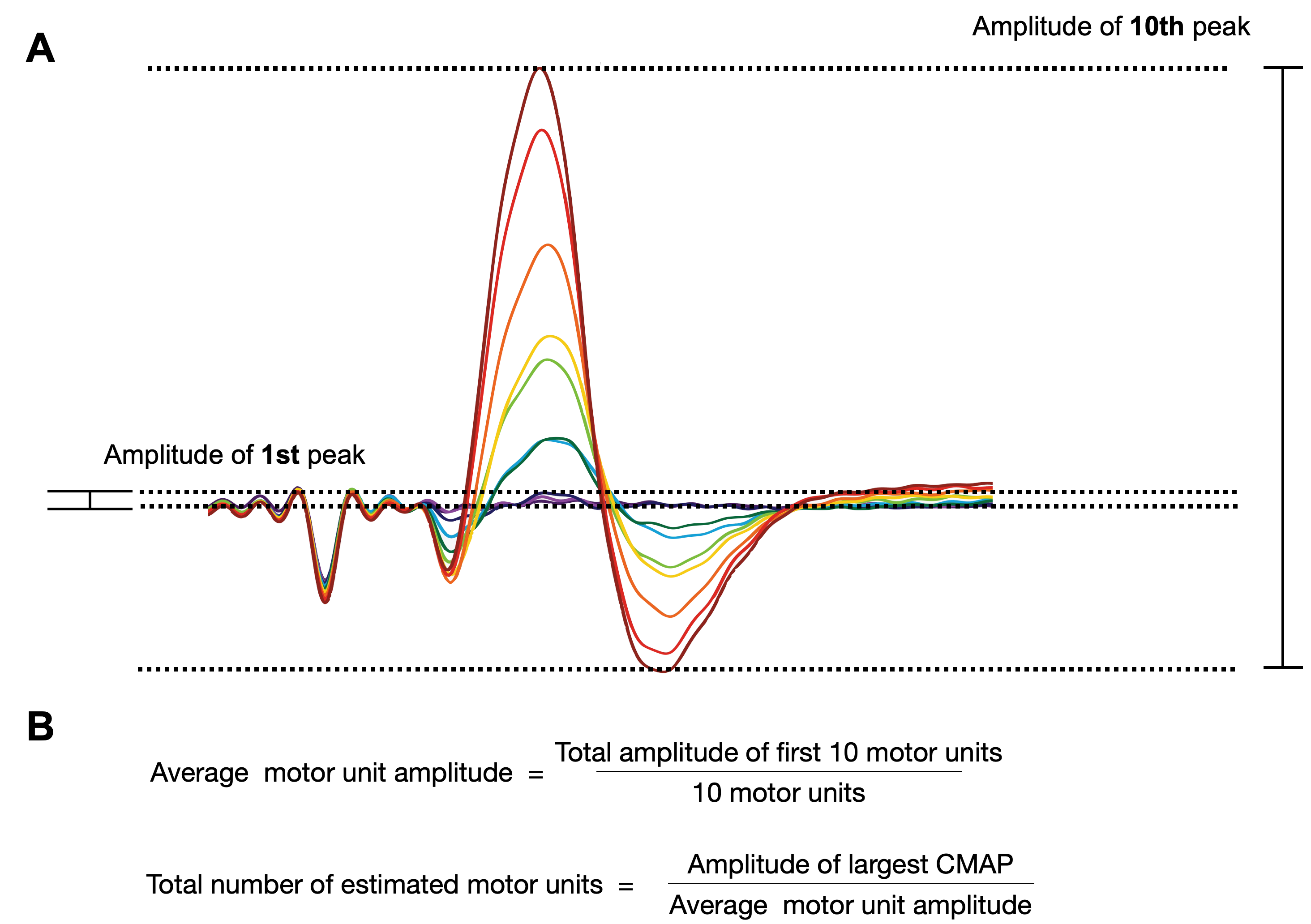

Figure S1.

(**A**) An example of CMAP traces recorded. For each current amplitude, 10 consecutive square wave pulses were applied. The average of the 10 evoked CMAPs were used. By increasing the current amplitude in small steps, we aimed to recruit *one* additional motor unit at every step (the first 10 steps are shown, each in a different color). The peak-to-peak amplitude of the first 10 peaks represents the combined amplitude of the first 10 motor units recruited. (**B**) We used the total amplitude of the first 10 motor units to find the average motor unit amplitude. Then, the maximum amplitude of all CMAPs was divided by the average motor unit amplitude to estimate the number of total motor units.

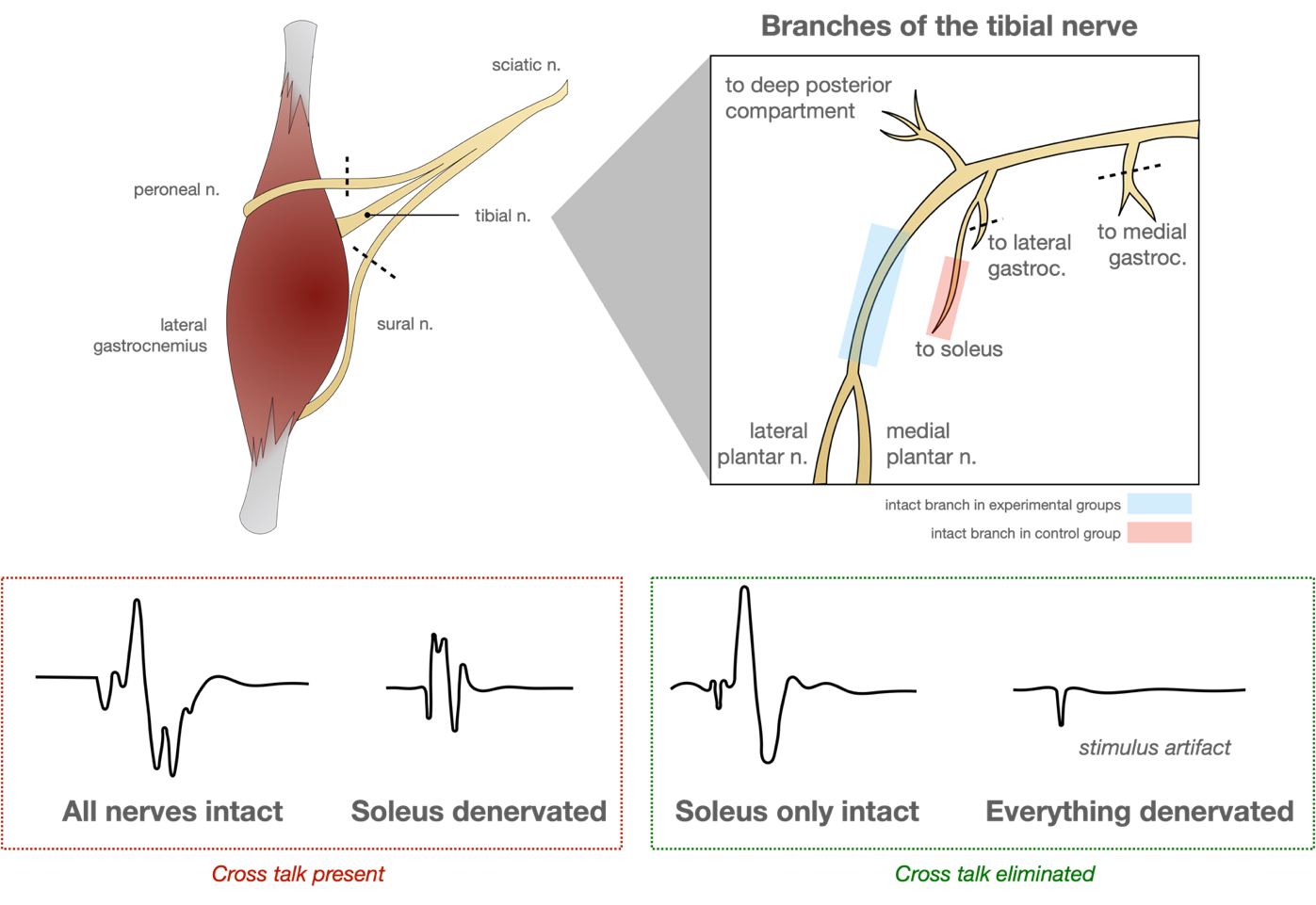
Figure S2.

To eliminate the risk of any cross talk from surrounding muscles affecting electrophysiology results, especially during electrical stimulation, we transected each sciatic nerve branch that did not directly innervate the muscle target, including the smaller branches of the tibial nerve. Each branch cut is designated by a dashed line. For experimental groups (i.e., reinnervated muscle), the only nerve branch left intact is shown in blue (the distal end of the tibial nerve that bifurcated into the lateral and medial plantar nerve. For the control group (i.e., intact muscle), the only branch left intact is shown in red (the original innervation of the soleus muscle). Evoked potentials demonstrate the presence of cross talk without (left, red) and with (right, green) this additional precaution of cutting other nerve branches.

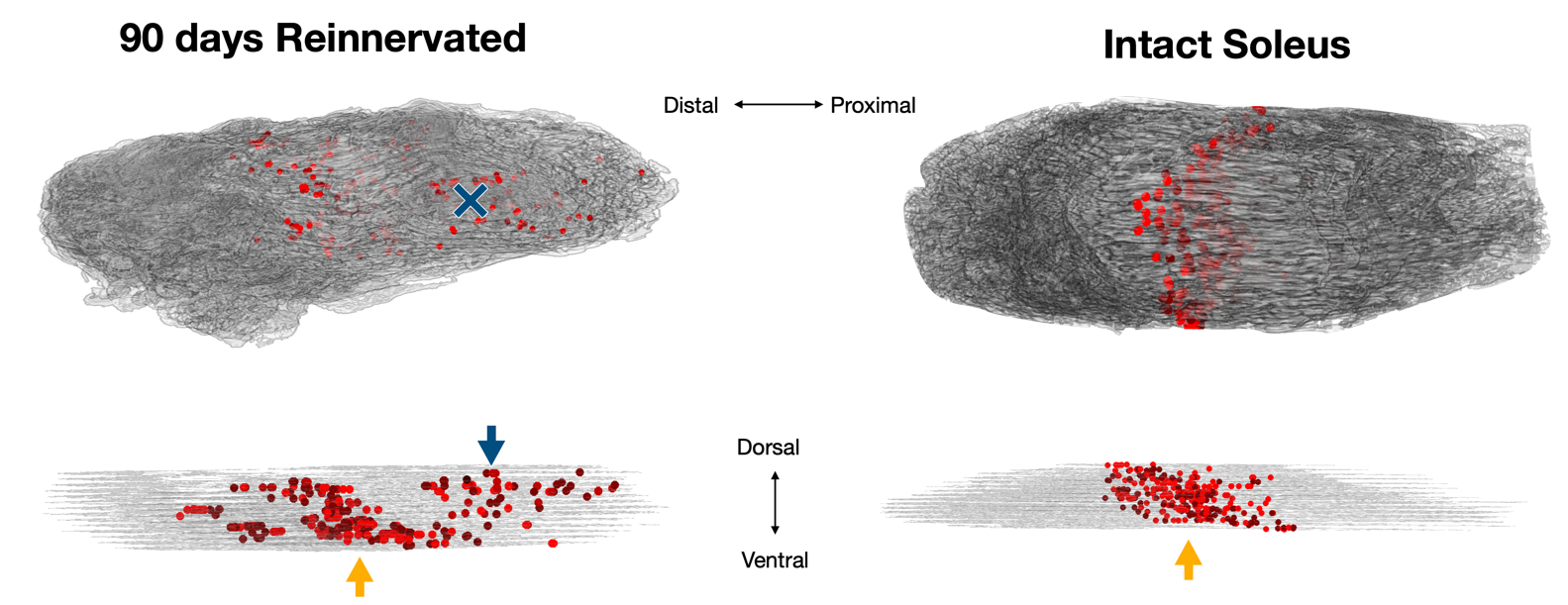
 Figure S3.

Representative images of reinnervated muscle 90 days post-surgery (left) and intact muscle (right) are shown. Each red dot represents a coordinate of an NMJ. The yellow arrow indicates the approximate original innervation point (located on the ventral side of the muscle). The blue arrow/blue ‘X’ indicates the approximate location where the donor nerve was transferred (located on the dorsal side of the muscle). Importantly, the 90-day muscle has ectopic NMJs near the transferred nerve site and thus exhibits a wider dispersion of NMJs compared to the intact control.

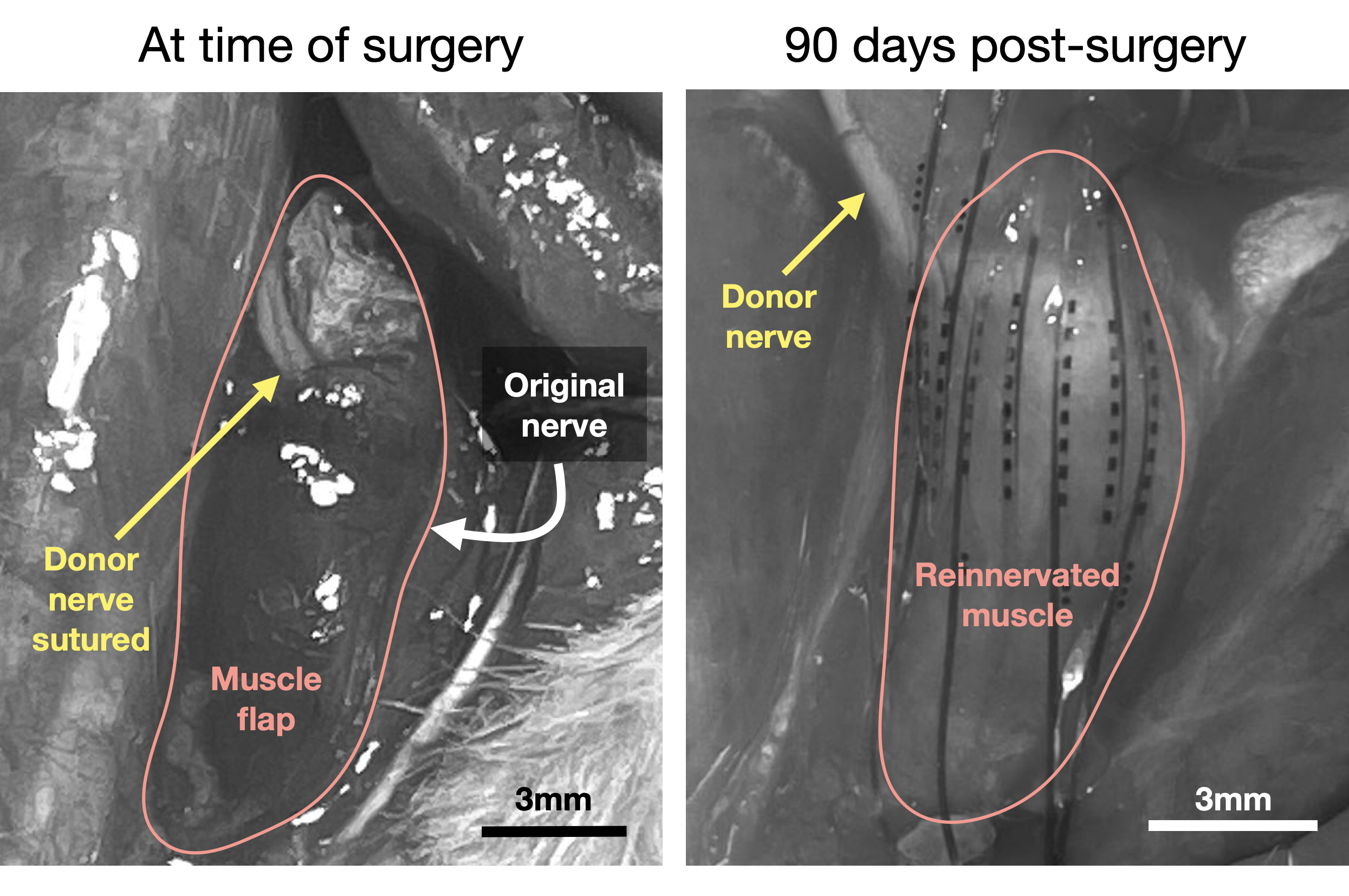
Figure S4.

During the initial muscle reinnervation surgery (left), the donor nerve, distal tibial nerve (yellow arrow), is sutured to the soleus muscle flap (outlined in red). Importantly, the muscle flap is denervated by cutting the original nerve (white arrow) to leave available motor endplates for the new donor nerve to reinnervate. Ninety days post-surgery, the reinnervated muscle is exposed (outlined in red). A 64-channel electrode array is placed on the surface of the muscle such that channels are located above and below the donor nerve (yellow arrow) as well as at the level of the original innervation point.

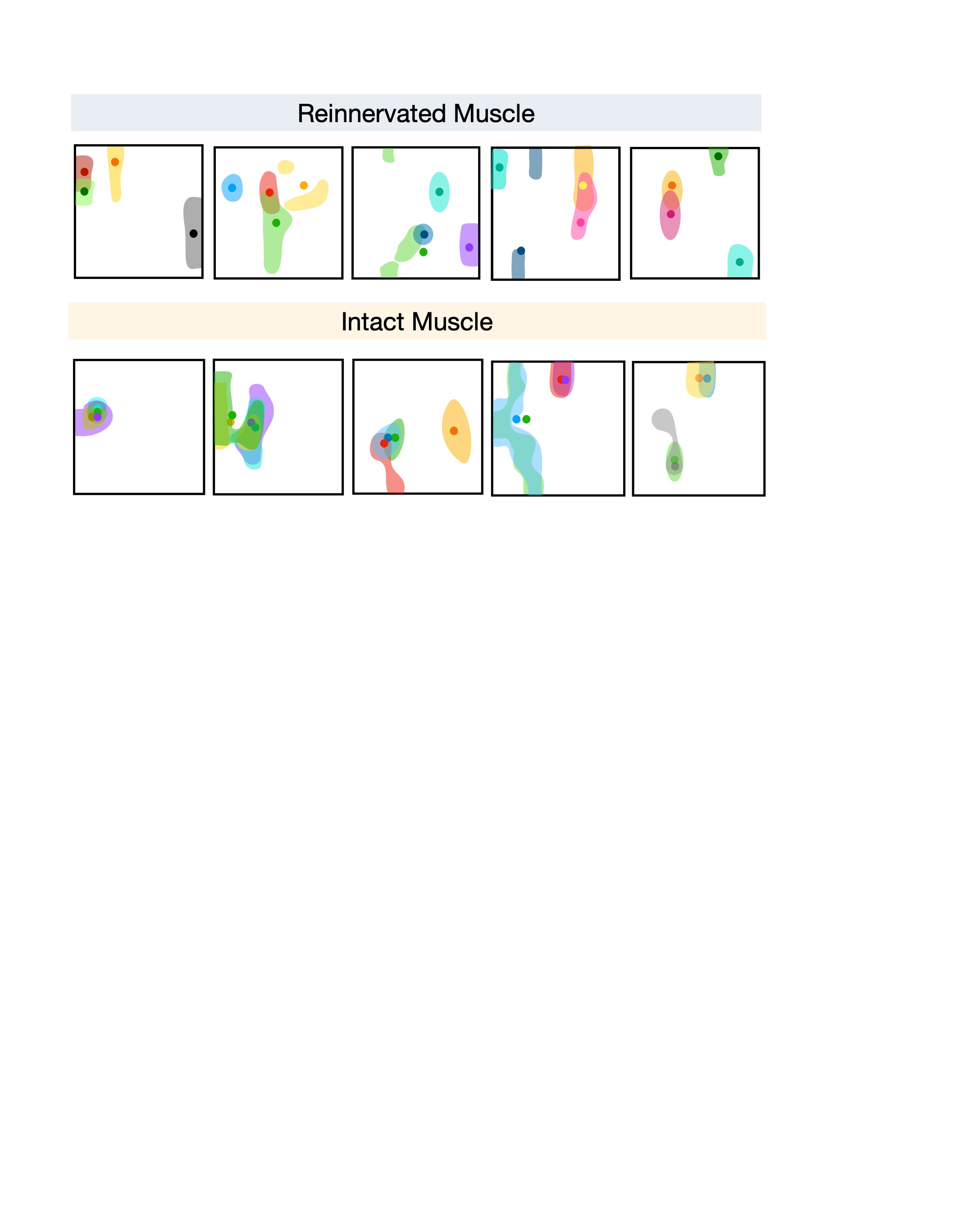
Figure S5.

Representative spatial maps in reinnervated (top) and intact (bottom) muscles. In each spatial map, four random motor unit territories are shaded in a unique color. The corresponding center of mass for each motor unit is designated by a point of the same color. More overlap between territories is seen in the intact muscle spatial maps.

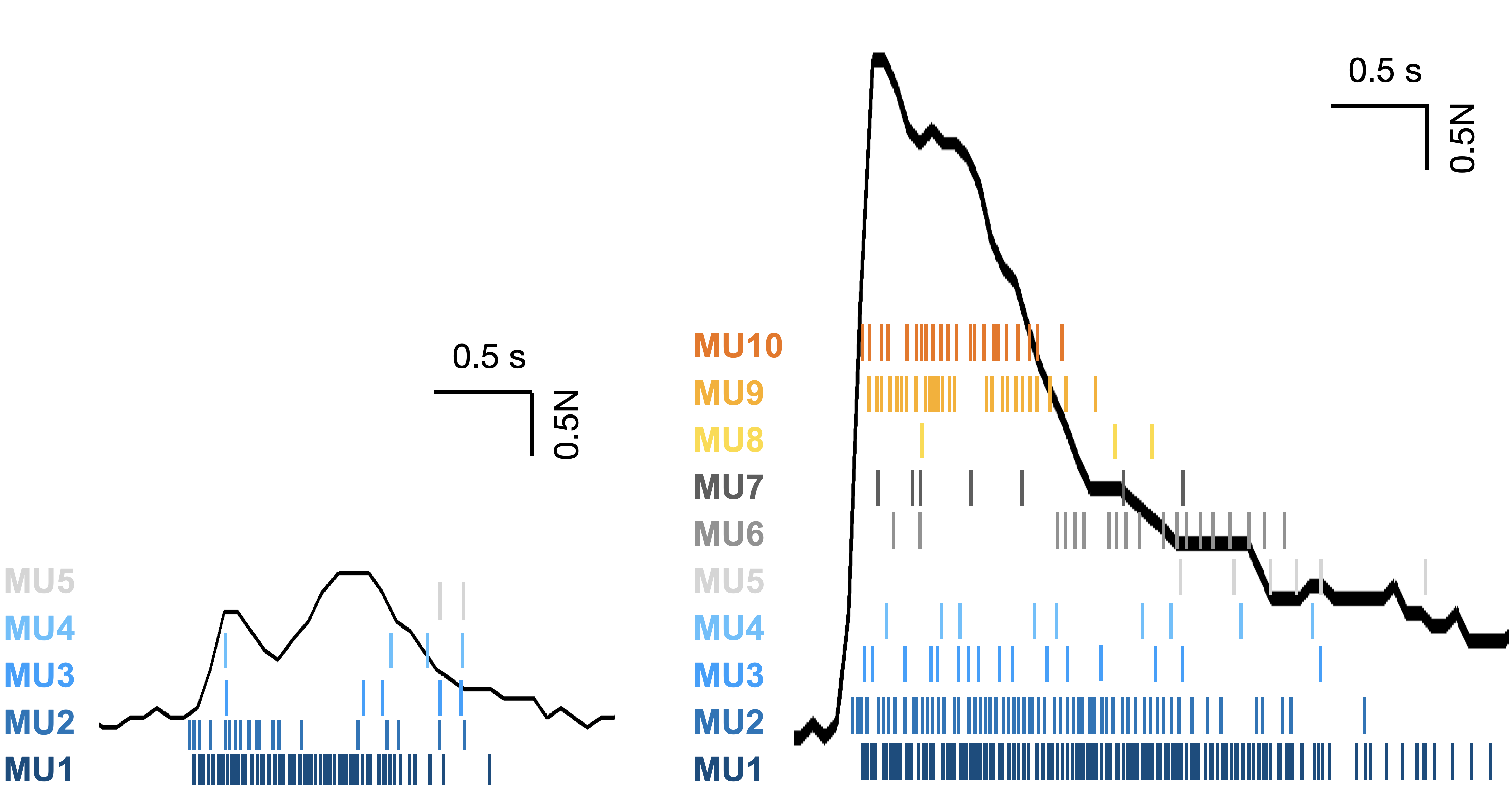

Figure S6.

Volitional motor unit activity and force production were recorded simultaneously. The force measured is depicted by the black curve. Two representative cases are shown, low force (left) and high force (right). In the reinnervated muscle flap, fewer motor units are active at lower forces. As force produced increases, the number of active motor units increases, and the firing rate of MU1-MU5 increases, just as they would in an intact muscle. Another notable point is that motor unit activity appears to mirror the subtleties in the low-force production case. The two ‘peaks’ are apparent in both the force production curve and motor unit activity.

| **Current range (μA)** | **Current step (μA)** |
| --- | --- |
| 20-120 | 2 |
| 125-300 | 5 |
| 310-500 | 10 |
| 500-750 | 50 |

Table S1.

Overview of electrical stimulation current amplitudes for evoking CMAPs.

| **Group** | **n** | **MUNE** | **Maximum CMAP Amplitude (mV)** |
| --- | --- | --- | --- |
| 15 days | 8 | 5.62 ± 5.32 | 0.20 ± 0.19 |
| 50 days | 7 | 12.17 ± 3.19 | 2.70 ± 3.44 |
| 90 days | 7 | 16.00 ± 4.83 | 3.46 ± 2.58 |
| Intact | 7 | 19.29 ± 8.90 | 6.80 ± 2.80 |

Table S2.

Overview of means and standard deviations for CMAPs and MUNE.

| **Parameter Name** | **Value** | **Explanation** |
| --- | --- | --- |
| freq_min | 250 | High pass cut off frequency |
| projection_threshold | 3, 3 | Waveform threshold, template threshold  Values help determine whether a detected spike belongs to a unit or is just noise (higher values result in stricter filtering) |
| minFR | 1/1000 | Minimum firing rate; Set very low for muscle recordings because spiking is expected to be more sparse than cortical data (especially using artificial stimulation) |
| sigmaMask | np.inf | Larger motor unit area |
| wave_length | 61 | Number of samples per waveform |
| nPCs | 3 | Number of PCs to use to build the embedding |

Table S3.

Overview of custom parameters used in Kilosort3 to modify the algorithm for motor unit sorting.

Movie S1.

Reinnervated soleus muscle target twitching in response to electrical stimulation. Surrounding muscles are silent because sciatic nerve branches were transected to eliminate crosstalk.

Movie S2.

Neuromuscular junction distributions in representative muscle 90 days post-reinnervation surgery.

Movie S3.

Live heatmaps of motor unit spatial activity during volitional plantar flexion for low (top) and high (bottom) force production.

Movie S4.

Virtual prosthesis controlled using motor unit activity (left) compared to the actual movement (right). Three representative cases are shown: low force, high force, and low force with a two-part flexion.
